# Submolecular-scale Hairpin DNA Folding Dynamics Studied by High-Speed AFM with Optical Tweezers

**DOI:** 10.1101/2024.08.14.605043

**Authors:** Kenichi Umeda, Shin’nosuke Yamanaka, Motonori Imamura, Fritz Nagae, Shingo Fukuda, Hiroki Watanabe, Takayuki Uchihashi, Shoji Takada, Toshio Ando

## Abstract

Optical tweezers have contributed to elucidate the folding mechanisms associated with biomolecules. By combining single-molecule or super-resolution techniques, imaging can also be performed while measuring or inducing force coupling with biochemical reactions; however, they cannot capture structural information beyond the fluorophore spatial resolution. To overcome this problem, here, we developed a technique that combines optical tweezers with high-speed atomic force microscopy (AFM). To solve the problem of incompatible instrumental configurations, we developed a unique optical tweezers measurement system that is specialized for high-speed AFM. Upon applying an external force to a synthesized DNA secondary structure, we successfully visualized the dissociation of the duplex structure. Furthermore, we succeeded in observing spontaneous reannealing of the duplex structure upon releasing the force, which demonstrates that the folding reaction can be reversibly controlled. We also reveal that along with duplex unfolding, a metastable secondary structure is generated and its topology changes transiently over time. The results indicate that this technique provides structural insights that cannot be obtained by conventional fluorescence techniques.

## INTRODUCTION

Optical tweezers (OT)^1-6^ is a powerful tool to evaluate the folding dynamics of biomolecules. This technique combined with single-molecule microscopy enables the manipulation of biomolecules while measuring the response to an external force simultaneously with imaging^6^. Thus far, it has been used to elucidate various biological processes including the hybridization mechanics of nucleic acids (DNA and RNA) ^4,6-13^, the formation of the cytoskeleton^14-16^, and the behavior of nucleotide-binding proteins. Despite significant interest in submolecular-scale structural changes induced by an external force, the spatial resolution of fluorescence microscopy remains limited by the size of the fluorescent label, which prevents direct observation at submolecular resolution. Although super-resolution microscopy has surpassed the diffraction limit, and provides further insight into detailed detail molecular structures^17,18^, spatial resolution is still constrained by the size of the fluorophore.

Over the past two decades, we have been developing high-speed atomic force microscopy (HS-AFM), which is a unique technique for probing the real-space dynamic behavior of biomolecules under physiological conditions at significantly greater spatiotemporal resolution compared with the fluorescence imaging (∼1 nm and ∼0.05 s)^19-28^. HS-AFM has revealed various biochemical functions stimulated by chemical^19^, photo^20^, or tip-induced-force^29^ methods. Folding researches have also been conducted using conventional AFM; however, they have not been able to observe the reversible dynamics of a structural change because the reactions are caused by chemical denaturation^30^ or meniscus force under ambient conditions^31,32^.

Unlike fluorescence microscopes, in HS-AFM experiments, the target molecules must be adsorbed onto a substrate, and the probe tip must scan across the surface, whereas OT normally measures a molecule tethered between two microparticles that are free from the surface interaction. This incompatible equipment configuration renders the combination of these techniques extremely challenging. In the present study, we established a combined system of HS-AFM and OT by devising the instrumental configuration. To realize synchronous control of these two techniques, we also developed a control software specialized for the HS-AFM/OT system. Furthermore, we developed a synthesis method for a long hairpin concatemer assay that is suitable for HS-AFM/OT experiments. Our system represents the first direct observation of a submolecular-scale duplex structural change including various transient states that have never been observed through conventional studies.

## EXPERIMENTS

OT traps a microparticle using an intensely focused laser beam through an objective lens. Manipulating the trapped microparticle enables a tensile force to be applied to biomolecules attached to the particles (Fig. 1 left). However, the OT optics cannot be incorporated under the sample stage of the standard sample scanning HS-AFM setup (Fig. 1 right). Therefore, we previously established a tip-scan HS-AFM (Fig. 1 center)^33^.

**Figure 1.**
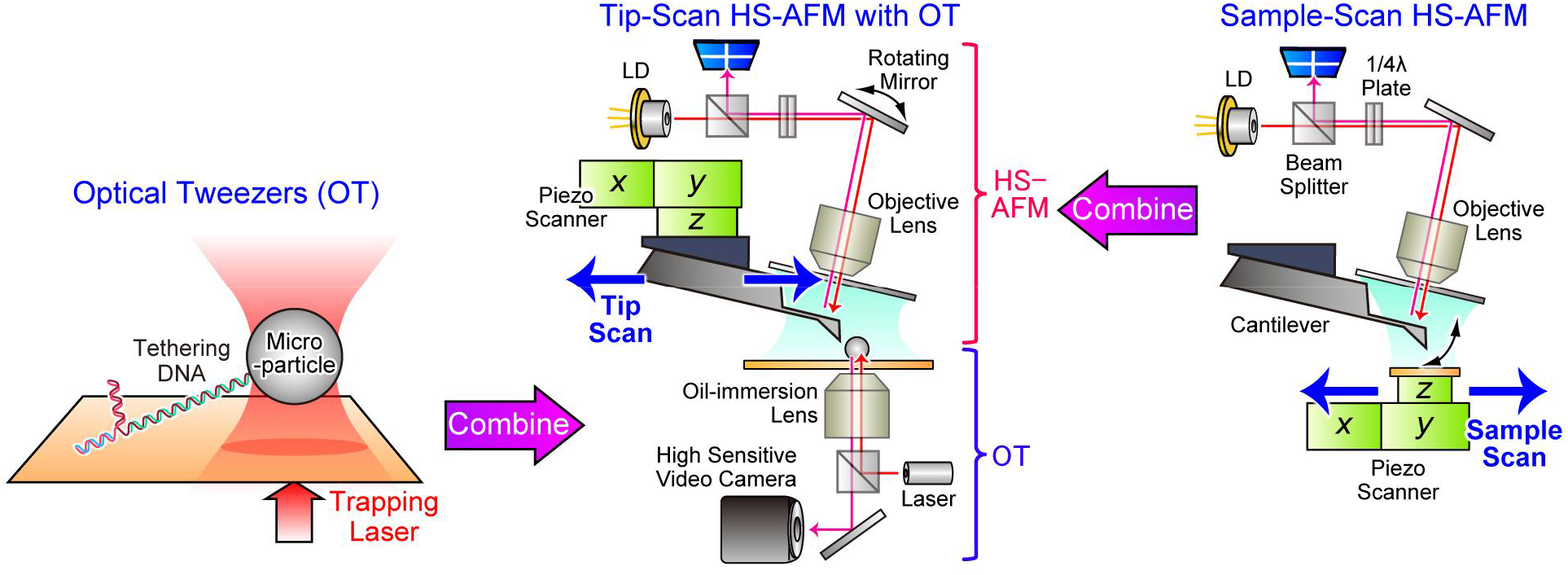
Combination of HS-AFM with OT. Schematics of OT (left), tip-scan HS-AFM-OT (center), and sample-scan HS-AFM (right).

To demonstrate the power of the system, we prepared hairpin DNA (also called stem-loop), a ubiquitous secondary structure formed from single-stranded DNA (ssDNA)^34^. It has also been used as a model system for studying the stability and mechanical properties of the duplex structure of double-stranded DNA (dsDNA) using single-molecule techniques^4,6-13^. To prepare this sample, we developed a synthesis strategy based on concatemer (a string of identical sequences) polymerization because incorporating multiple hairpins into a single DNA with several microns in length facilitates the location of hairpin structures during AFM measurements (Supplementary Note 1).

We evaluated the structure of the hairpin concatemer using HS-AFM. A DNA molecule with a length of 19 μm (Fig. 2a) was synthesized, in which five hairpin constructs were incorporated (Fig. 2b). By measuring the intervals of the hairpin structures, we confirmed that it had a multiple of 0.9 μm. The individual hairpins consisted of a 93-bp stem with 55.9% GC content that was capped by a (dT)_6_ loop and flanked by two handles through (dT)_3_ ssDNA (Fig. 2c). Thus, the intact hairpin stem length was ∼32 nm (0.34 nm/bp), whereas the unpeeled ssDNA length was predicted to ∼67 nm (Fig. 2d).

**Figure 2.**
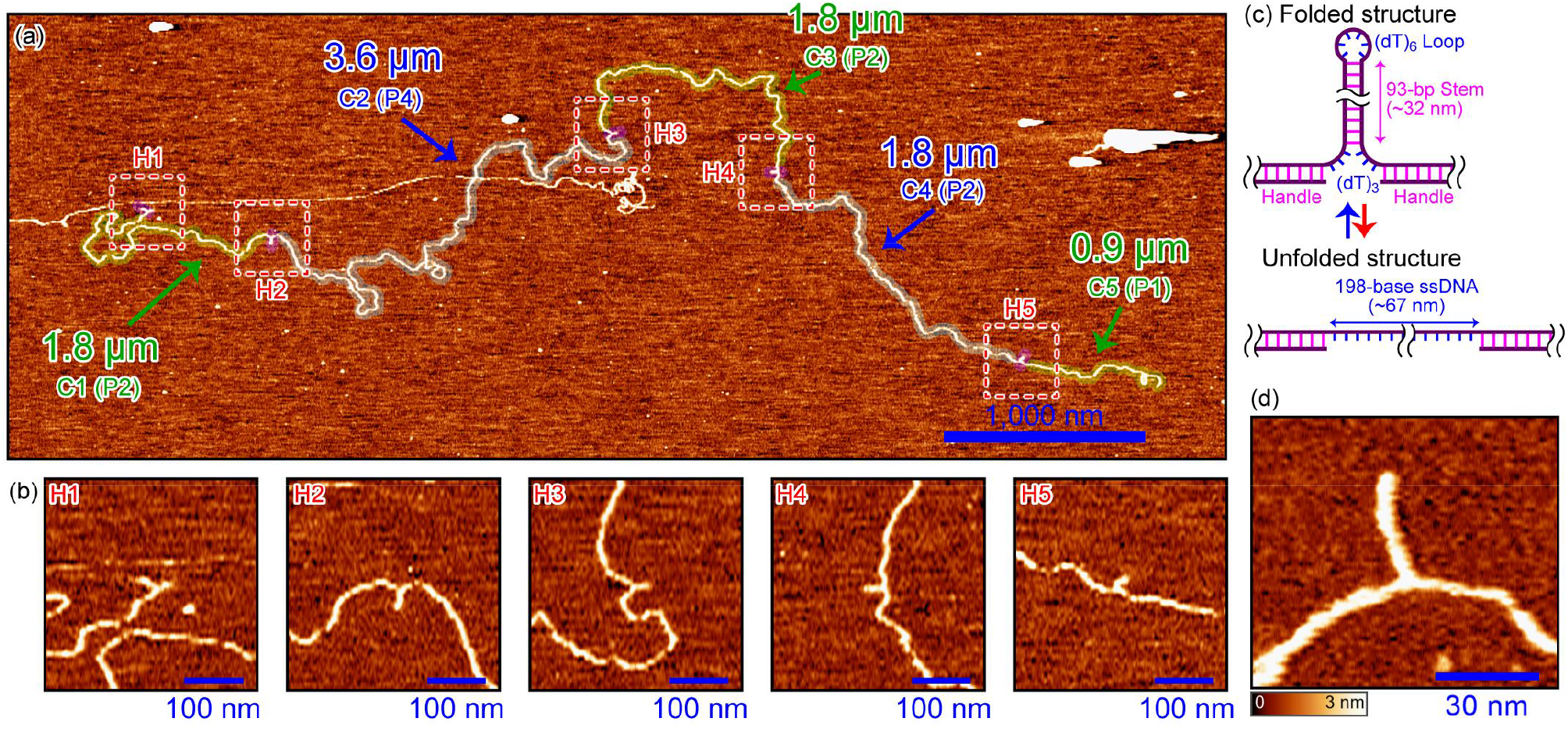
Design of the hairpin DNA concatemer assay. **a**, Overall HS-AFM image of the hairpin concatemer. C1–C5 and H1–H5 indicate the serial number of the linear regions and hairpin structures, respectively. The number shown in brackets represents the number of linearized plasmid DNA molecules contained in each linear region, which can be calculated by dividing the contour length by one unit length of ∼0.9 μm. **b**, Close-up images of each hairpin structure segmented from the overall image. **c**, Schematics of the synthesized hairpin constructs with the folded (**a**) and unfolded (**b**) structures. **d**, High-resolution image of a hairpin structure.

For normal OT measurements, both ends of the DNA are attached to particles and tension is applied to the DNA using two particles. However, since HS-AFM cannot visualize molecules in a floating state, only one end of the DNA molecule was biotinylated for coupling to a streptavidin-functionalized microparticle (Supplementary Note 2), and the other end was adsorbed to the surface. Because DNA is not able to move when strongly adsorbed to a substrate, it was electrostatically weakly adsorbed to the substrate through dissolved electrolytes. We used muscovite mica as the substrate because coverslips commonly used in OT measurements has a rough surface that prevents DNA from freely diffusing. When mica is thinly sliced, it becomes transparent and allows the laser to pass through. Because DNA was weakly adsorbed, it freely diffused across the surface when no external force was applied using OT (Supplementary Fig. 5, Movie 2). These conditions are essential for observing the spontaneous occurrence of the duplex reannealing process as discussed later.

In general, OT experiments are performed with bovine serum albumin (BSA) solutions to prevent particles from adsorbing to the substrate; however, since HS-AFM requires a clean substrate, BSA cannot be used. Therefore, we used 4.41-μm particles, which are larger than the 1–2 μm particles used in common OT to obtain large radiation pressure. This made it possible to re-trap particles even stuck to the surface.

## RESULTS

### Optical tweezers experiment

In our optical setup, the cantilever and microparticle were located on the left and right sides, respectively (Fig. 3a,b). In the first experiment, we performed HS-AFM imaging while moving the particle to the right at a constant velocity of 138 nm/s until hairpin unfolding was observed. Next, we moved it back to the left at the same velocity (Fig. 3a). For normal OT measurements, the microparticle position is measured by detecting scattered light using a segmented photodiode on the top. However, this method cannot be used for the HS-AFM/OT system because of the AFM scanner and optics. Therefore, the particle analysis was performed using image processing based on a sub-pixel-resolution cross-correlation technique (see the methods for details). We quantitatively analyzed the particle trajectory and force–extension curve from the optical images (Fig. 3b), which showed the stick-slip movement of the particle from one adsorbed site to another (Fig. 3c,d). This indicates that the force estimated from the particle motion includes the effects of the surface interactions as well as the force exerted on the molecule. Although the forces applied to the hairpin can be roughly estimated by analytical calculation as discussed later, it is preferable to further increase the laser power to reduce the surface effect.

**Figure 3.**
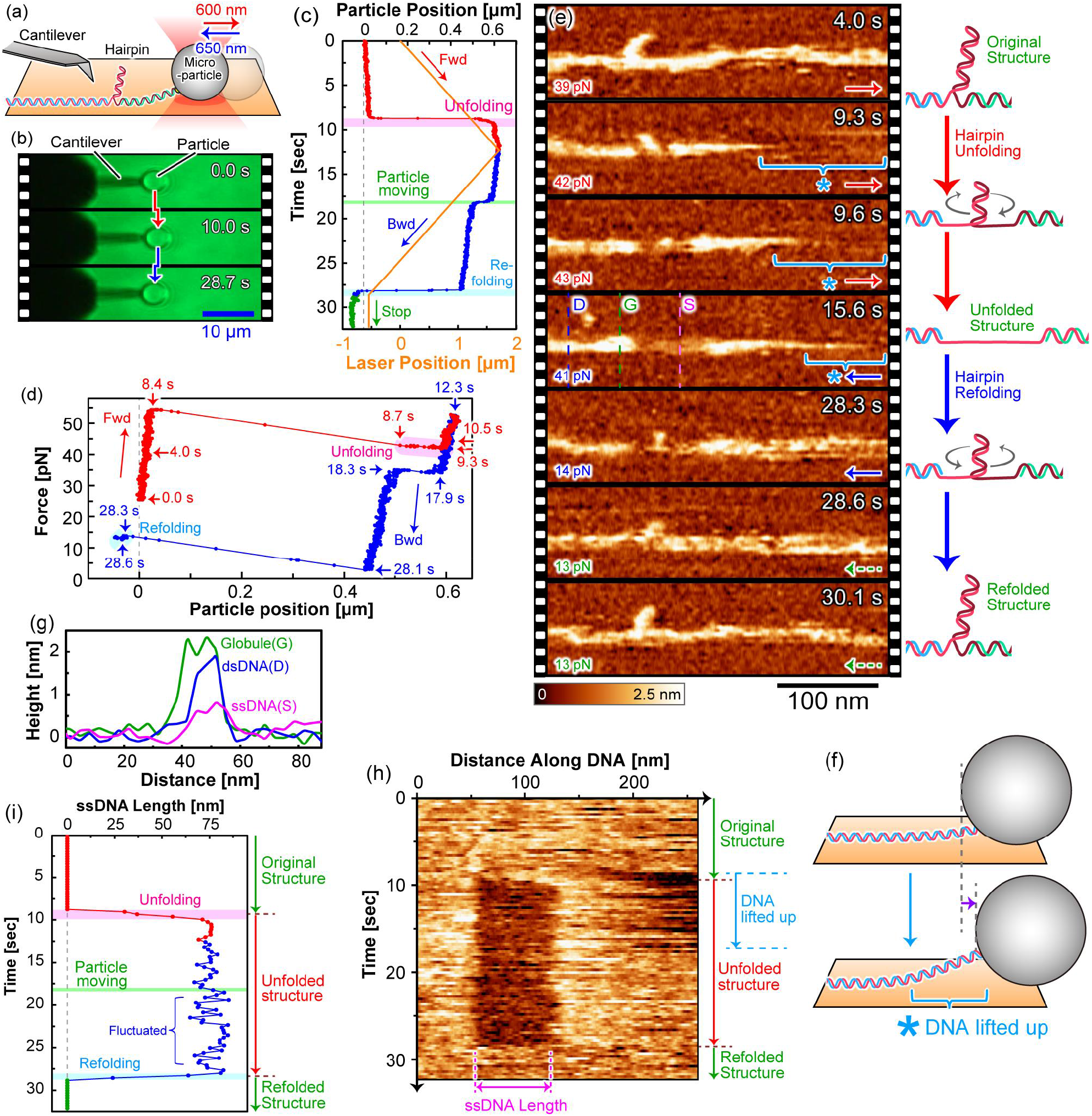
Hairpin dynamics with right conditions. **a,b**, Schematic of the experiment (**a**) and time-lapse optical video microscopy images (**b**). **c,d**, Particle trajectory (**c**), and converted force curve (**d**). In (c), the orange line in the inset indicates the trapping laser trajectory. In (c,d), the red, blue, and green lines indicate the trapping laser in the forward, backward, and stop conditions, respectively, and the pink and cyan regions indicate the data points where the unfolding and refolding events were observed in the HS-AFM images, respectively. **e**, Time-lapse HS-AFM images (left) and the schematics of the hairpin structures (right), where the rotating arrows indicate the torsional force direction accompanied by the hairpin folding events. The original data is shown in Supplementary Movie 4. **f**, Schematic of DNA lift event, which is indicated by an asterisk at 9.3–15.6 s in Fig. 3e. **g**, Line profiles extracted from Fig. 3e at the position indicated by the broken lines of dsDNA (D), globule (G), and ssDNA (S). **h,i**, Kymograph of the molecular height along the DNA ridge line (g), and time course of unpeeled ssDNA length estimated from the kymograph (h).

Note that there is the possibility that another hairpin was inserted in the tethered DNA because the length of the tethered DNA was estimated to approximately 1.8 μm based on the geometry of the cantilever and particle (Supplementary Fig. 6 and Supplementary Note 2). However, if a hairpin was also included in the tethered region, a force plateau would appear at 15 pN, corresponding to the intrinsic hairpin unfolding event^9,12^, because the tethered DNA is free from the substrate. The force curve data did not exhibit a force plateau (Fig. 3d), suggesting that the hairpin was not incorporated in the tethered DNA region.

### HS-AFM imaging of hairpin dynamics

In Fig. 3e, the HS-AFM image at 4.0 s shows the hairpin structure with a ∼35-nm length as designed. The particle remained stationary for the first 8.4 s of observation (Fig. 3d). Subsequently, a rapid 500 nm displacement occurred (8.4–8.7 s), followed by a gradual 100 nm movement over several hundred milliseconds (8.7–9.3 s). Concomitant with this movement, the right portion of the DNA disappeared, indicating that it was lifted up from the surface by the particle force (Fig. 3f). This was unexpected because we anticipated that DNA would be difficult to observe because the probe tip would get stuck in the DNA when it was lifted. We found that the AFM tip had a minimal impact on the tethered DNA only when the tip scanning direction and the pulling direction were the same (Supplementary Fig. 7, Movie 3). During the gradual movement of the microparticle, the hairpin structure was unfolded to expose the ssDNA region over 0.9 s (8.4–9.3 s). A decrease in the DNA height from 2 to 0.4 nm rendered the ssDNA and dsDNA regions clear (Fig. 3g). Finally, the total ssDNA length was extended to ∼70 nm, which was double the hairpin length. We depicted the schematics on the right side of Fig. 3e based on molecular dynamics (MD) simulation results, which showed that the unfolding progresses while the hairpin stem rotates. The force exerted on the particle reached 55 pN, which was significantly greater than 15 pN. We analyzed the mechanics using a simple viscoelastic model, which revealed that most of the force was used for moving the particle and only 21 pN was exerted on the DNA (Supplementary Note 3).

Kymograph analysis (Fig. 3h,i) revealed that after gradual movement, the ssDNA length and the microparticle position were unchanged, which indicated that the force exerted on the DNA was not strong enough to break the dsDNA (10–28 s). Moving the particle back to the left resulted in the immediate re-attachment of the lifted portion of the DNA (17 s). Following this movement, the ssDNA length fluctuated more, indicating a weakened tensile force. Subsequently, the ssDNA refolded to its original hairpin structure within 0.9 s (28.0–28.9 s), which was identical to the unfolding time.

If the tethered DNA is free from surface interactions, its unfolding and refolding transient time is estimated to be ∼84 μs based on the previously reported value of ∼0.9 μs/bp^35^. This is 12,000-fold faster than the 0.9 s observed in the HS-AFM measurement, which may be the result of surface interactions. The unfolding and refolding are based on different mechanisms. The unfolding is induced by the tensile force from the particle pulling, whereas the refolding is a spontaneous event driven by hydrogen bond formation between the complementary base pairs. Therefore, if the pulling force significantly exceeds the refolding force, the unfolding time would be shorter compared with the refolding time. However, because the hairpin was pulled with a force of approximately 21 pN, which is close to 15 pN, it was in a near-equilibrium state^9,36^ and the transition times were comparable.

Previous studies using OT demonstrated that an intermediate state is observed by maintaining the equilibrium condition by controlling an external force^9,36^. To determine whether an intermediate state exists also in our data, we analyzed the unfolding data in more detail. First, the particle trajectory in Fig. 4a exhibited two plateaus while in the ssDNA length in the HS-AFM data (Fig. 4b) exhibited only one plateau. Considering the hairpin structure in the HS-AFM data in Fig. 4c, we assumed that the first plateau at 8.8 s corresponded to a state in which the tethered DNA was taut, and the second plateau at 9.1 s is a transient state of unfolding (Fig. 4d). The plateau at 9.2 sec in the ssDNA length corresponded to the second plateau of the particle movement, which indicates the existence of a metastable state during unfolding.

**Figure 4.**
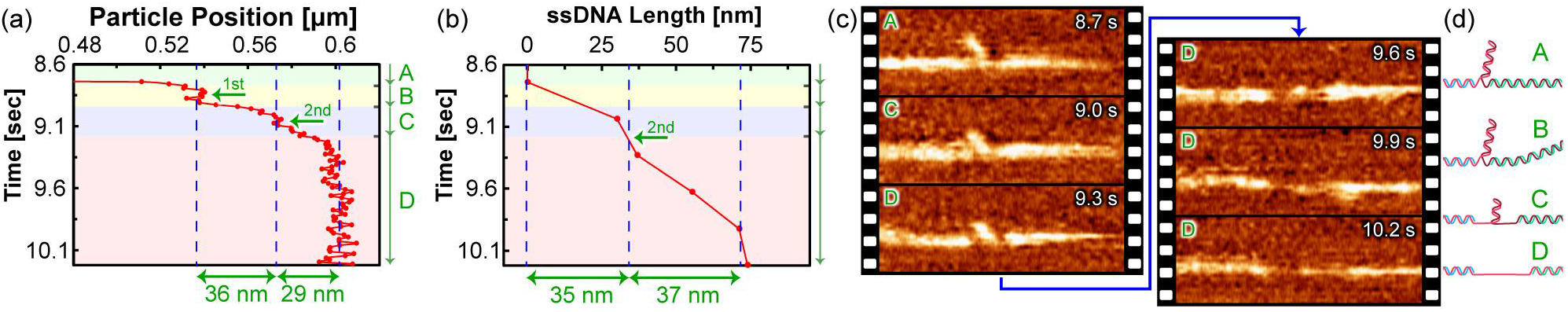
Detail analysis of the unfolding event. **a,b**, Zoom-in curves of the particle position (a) and ssDNA length (b) during the unfolding event. The original data are shown in Fig. 3(c) and 3(i), respectively. **c,d**, Close-up AFM images of the hairpin construct during the unfolding event with the corresponding state indicated at the top left (c) and Schematics of the hairpin structures in different unfolding states (d).

### MD simulation of hairpin dynamics

To examine the origin of this metastable region, we performed an MD simulation^34,35^ that explicitly takes the surface effects into account (Supplementary Note 4) and successfully reproduced the reversible folding reactions of the hairpin structure (Fig. 5a). From the structural data, we generated pseudo-AFM images, which highlighted the transient formation of small secondary structures (globules indicated by arrows in Fig. 5b) during the refolding process. This originates from the transient rupture and recombination of the hydrogen bonding while searching for a more energetically favorable structure^36^. In principle, this phenomenon may also occur during unfolding, but it was not observed in the simulation because we applied an external force larger than the hairpin annealing force to reduce the simulation time, which renders it a non-equilibrium process. The metastable structure observed in the experiment is possibly the result of this transient hydrogen-bond search reaction. In experiments, this phenomenon was observed only during unfolding, contrary to the calculations, but if the tensile force by the particles exactly balances the hairpin force, it may also be observed during the refolding process.

**Figure 5.**
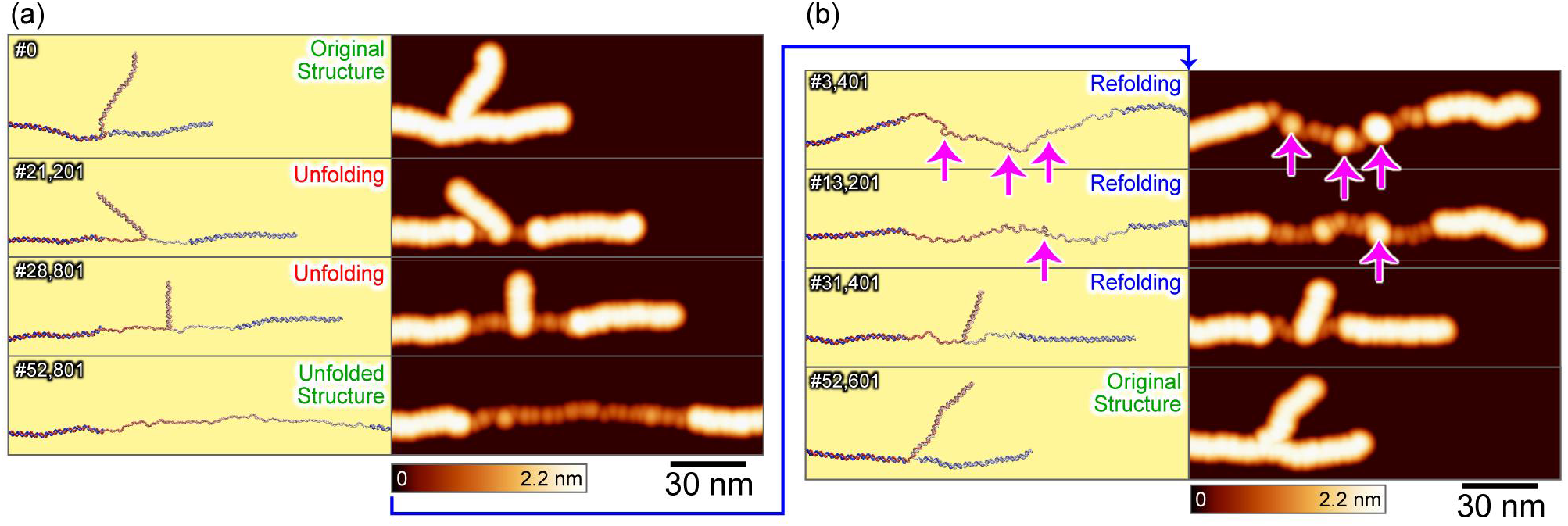
MD simulation of hairpin unfolding and refolding. **a,b**, Snapshots (left) and pseudo-AFM images (right) of MD simulation during the unfolding (**a**) and refolding (**b**) processes. The original data are shown in Supplementary Movie 5.

### HS-AFM imaging of hairpin overstretching

In the second experiment, we pulled the microparticle further away even after hairpin unfolding was complete (Fig. 6a,b). The particle trajectory and force curve exhibited a greater number of stick-slip motion of the particle compared with that in Fig. 3 (Fig. 6c,d). In Fig. 6e, the first HS-AFM image (6.1 s) shows the intact hairpin structure. After 45 pN was exerted on the particle, the first particle jump was observed, which stimulated the hairpin unfolding (7.3 s). The unfolding was complete after one second as with the previous experiment. At this stage, the full-length ssDNA was ∼70 nm (10.8 s). Then the particle was further moved toward the right (17.2–21.2 s) and eventually reached a 1,300 nm distance from the original position.

**Figure 6.**
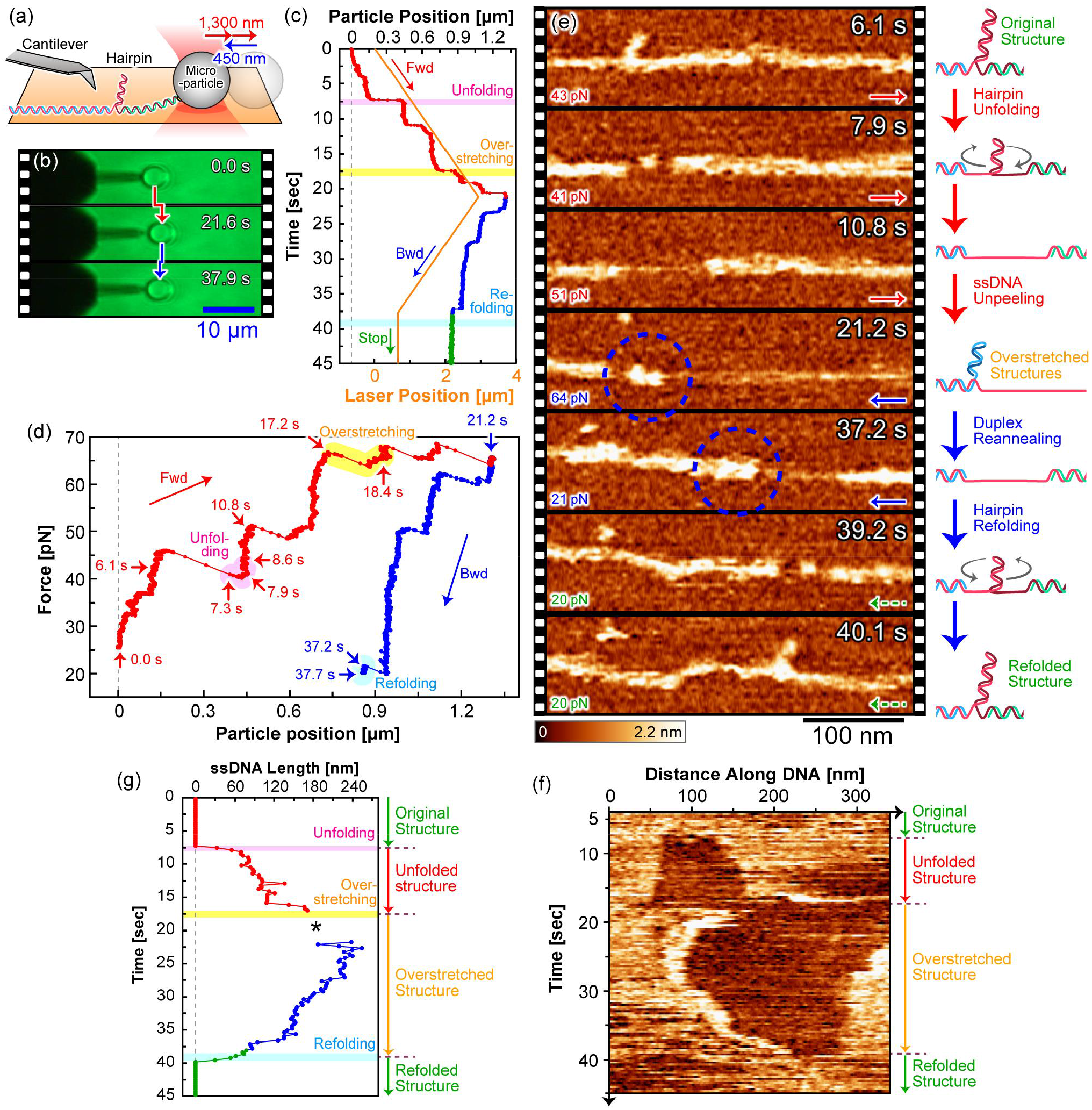
Hairpin dynamics under overstretching conditions. The descriptions of Fig. 6a–e and Fig. 6f,g are the same as those of Fig. 3a–e and 3h,i, respectively. The original data may be viewd in Supplementary Movie 6. In Fig. 6g, the asterisk indicates the region where the ssDNA length cannot be quantitatively estimated because the ssDNA length exceeds 240 nm and the right boundary of the ssDNA is outside of the image.

Kymograph analysis revealed the extent of the ssDNA region further extended to ∼200 nm along with this particle movement (Fig. 6f,g). This indicates the occurrence of the overstretching transition, in which the ssDNA was unpeeled from the discontinuity between the hairpin stem and handle of the dsDNA. The unpeeling force applied in this study was 65 pN, which is the same as the force estimated in the previous OT studies^7^. Unlike the hairpin unfolding event, the intrinsic transition force was observed because it was transiently lifted and free from the surface interaction (Supplementary Movie 5).

We subsequently moved the particle back to the left by 450 nm. Following this movement, the left part of the DNA was gradually reannealed to recover the duplex structure, of which the mechanical stress biased the DNA diffusion to the right. The right portion of DNA was not moved because it was still weakly strained by the particle. Kymograph analysis (Fig. 6f,g) revealed that the ssDNA length was gradually reduced from 240 nm to 70 nm (21.2–37.2 s). The unpeeling was completely recovered, immediately followed by the refolding of the hairpin structure over 0.9 s (39.2–40.1 s). Interestingly, the duplex structure was completely reannealed even after the secondary structures were formed from unpeeled ssDNA.

In Fig. 6e, because one or two globules were nucleated from the boundary between the ssDNA and dsDNA (designated 1-globule and 2-globules states, respectively), we retrieved the close-up images in Fig. 7a. The original structure exposed only the 1-globule state (18.8 sec), but the size significantly increased at 20.3 s. Then the number of globules increased to two at 21.5 s, but they were merged into one globule again at approximately 26 s. Interestingly, the globule was split into two small globules at 36.9 s and eventually disrupted at 38.7 s. To determine the changes in the globule state, we quantified the time course of the number and size (volume) of the right end of the globule in Fig. 7b. The results indicated that the number of globules changed every 2–3 s and the apparent size of the globules was positively correlated with the number of globules. Therefore, in Fig. 7c, we quantified the globule volume for each state and found that 2-globules had approximately half the volume of 1-globule.

**Figure 7.**
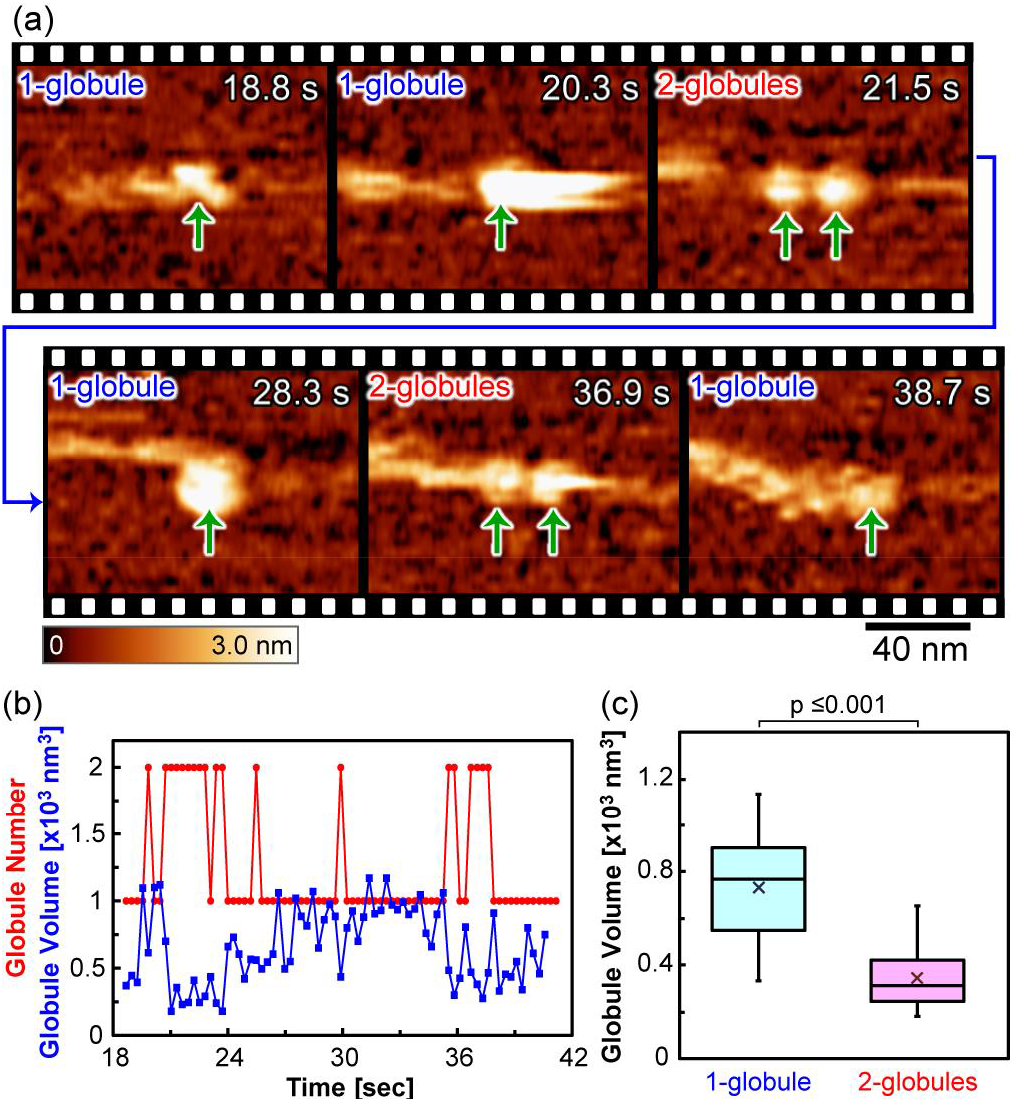
Secondary structure nucleated from unpeeled ssDNA. **a**, Time-lapse HS-AFM image of the boundary between dsDNA and ssDNA, cropped from the blue broken circles in Fig. 5(e). The original data may be seen in Supplementary Movie 7. **b,c**, Time course of the number and volume of globule (b), and quantitation of the globule volume in 1-globule and 2-globules states. Statistical differences were examined by the Welch’s t-test (c).

This globule was most likely a transient secondary structure as observed in Fig. 5 because the unpeeled ssDNA cannot form a complementary dsDNA, rather it forms a stable secondary structure. We also predicted a stable secondary structure by simulation, which revealed multiple stable structures (Supplementary Fig. 10). Therefore, the morphological change observed in AFM probably reflects the topological change of the secondary structures.

We did not expect that duplex folding could also be observed even under the surface interaction. It was not trivial that the original structure could be completely recovered without misfolding regardless of an estimate of the numerous local minima in the energy landscape resulting from the adsorbed condition. To identify the molecular-scale mechanism of the unpeeling reaction, we performed an MD simulation of the overstretching transition and reproduced the spontaneous recovery of the original dsDNA structure even under the surface interaction (Supplementary Fig. 11, Movie 8).

## Summary

In summary, we succeeded in developing an HS-AFM system in combination with an OT apparatus, which revealed the submolecular-scale unfolding dynamics of hairpin DNA stimulated by an external force and refolding dynamics, which spontaneously arise along with the release of the external force. Two of the transient times were approximately 1 second, 12,000-fold slower than the intrinsic ones due to the diffusion-limited conditions. During the overstretching condition, we also captured the denaturation of the duplex structure and the reannealing processes. There is still ample room for further improvement in this system; however, the present system will benefit numerous applications in biological research such as protein folding and DNA binding proteins (polymerases, helicases, etc.). It will constitute a core technique for biological and pharmaceutical studies.

## Methods

### 1. Synthesis of Hairpin Concatemer

To observe the hairpin folding dynamics using HS-AFM, we developed a hairpin concatemer synthesis and assay (Supplementary Fig. 1). The DNA sequences used in this study are listed in Supplementary Table 1. We first constructed three small dsDNAs, hairpin root, hairpin cap, and biotin end from oligo ssDNA synthesized by PCR Primer suppliers (IDT and Eurofins) through an annealing process. We used a pUC19 vector (2,686 bp) from New England Biolabs (NEB) because it has single BsaI restriction endonuclease site, which was cleaved to create ligation overhangs outside of the recognition site with a non-palindromic sequence. An aliquot of pUC19 was digested by BsaI-HFv2 (NEB, R3733L) at 37°C for 16 h. After digestion, for subsequent ligation, the remaining enzyme was inactivated at 80°C for 20 min. There are three BsaI recognition sites in the hairpin root, each of which has a different cleavage sequence. The ligation overhang sequence in the hairpin loop was complementary to that in the hairpin cap. One ligation overhang sequence in the handles was complementary to another one, which was equivalent to the cleavage sequence in pUC19. Finally, these constructs were mixed and ligated to produce long-chain concatemers with a random insertion. The number of inserted hairpins depends on the mixture ratio between pUC19 and the hairpin root. If the hairpin root fraction increases, the hairpin insertion increases, but the length of the concatemer decreases due to the incomplete-length oligonucleotides, which leads to a poor yield. Since the product contains concatemers with various lengths, we purified only long concatemers using electrophoresis in 0.3wt% PrimeGel Agarose GOLD 3-40K (TaKaRa Bio).

### 2. Optical tweezers system

For the OT measurements, a micromanipulation system (SIGMA KOKI, MMS-1064-2000-2L/2E/2S) was used with a 1,064 nm Nd:YAG Laser at a maximum power of 2.0 W, which was controlled by a dual-axis mirror galvanometer (Supplementary Fig. 3). As a substrate, we used muscovite mica (Furuuchi Chemical Co. Ltd.) with dimensions of 25 × 25 × 0.1 mm^3^. To focus the laser beam onto the substrate, a 100× objective lens (NA 1.49, Nikon, CFI SR HP Apochromat TIRF 100XC Oil) was used with immersion oil (Olympus, IMMOIL-500CC). The refractive index was 1.518, just slightly smaller than 1.55–1.61 of mica. The bright field image of the sample surface was illuminated with a fiber light through the overlaying optics for HS-AFM and monitored by a high-sensitive color video camera (Canon, ME20F-SH), which had a 35 mm CMOS sensor with 1920 × 1080 pixels^2^ at 60 fps. BrightLine basic 530/43 Fluorescence Filters (Semrock Inc) were used for eliminating the 780-nm laser beam (Rohm, RLD78MZA6) used in the AFM optical beam deflection system to analyze the particle trajectory. Although the pixel resolution was 88 nm/pixel, the net spatial resolution was improved to less than 10 nm/pixel using a cross-correlation algorithm with parabola fitting. To facilitate particle manipulation, we developed software that controls the laser position and shutter using a wireless PlayStation4 gamepad (Sony Interactive Entertainment) through an RS-232C serial communication protocol (Supplementary Fig. 4a). We used a streptavidin-functionalized polystyrene microparticle suspension (Spherotech, SVP-40-5) with an average diameter of 4.42 ± 0.3 μm. In this optimized system, the particle can be freely manipulated, even after it is stuck to the surface. We demonstrated the manipulation system by drawing characters (Supplementary Fig. 4b).

### 3. Optical tweezers experimental condition

We prepared a hairpin DNA stock solution of 0.5 ng/μL concentration, which was further diluted 8-fold with 10 mM Tris-HCl (pH 7.5) prior to each experiment. An 8 μL aliquot of a particle suspension, vortexed beforehand, was slowly added to an 8 μL aliquot of this diluted solution while stirring using a pipette tip. The final concentration of the particle was 0.177 pM, whereas that in DNA assay was 30.5 pM assuming that the DNA length was 25 kbp. We optimized the particle/DNA mix ratio to approximately 1:20. Next, the DNA-particle-conjugated solution was incubated in a refrigerator for at least 1 h. The solution was stirred with a pipette tip three times during the incubation.

Normally, a transparent coverslip is used as a substrate in OT experiments, but an atomically flat and non-contaminated surface (e.g., mica) is preferable for the bioAFM experiments. The mica is also a candidate for the OT substrate because it becomes transparent if it is cleaved into as thin as 0.05 mm thickness. We first sliced the mica in half using a cutting knife, and then further cleaved the intact face using Scotch tape. For sealing a liquid droplet, a Teflon sheet (22 × 22 × 0.5 mm^3^ with a φ7-mm center hole) was glued onto the mica surface using an epoxy resin. After the adhesive cured for 10 min, a 1 μL aliquot of the incubated solution mixed with 38 μL aliquot of 100 mM NiCl_2_ (≧98.0%, Nacalai Tesque) in 20 mM Tris-HCl (pH 7.5) was dropped onto the surface and incubated for 4 min. Then, it was gently rinsed with a solution containing 30 mM KCl (Nacalai Tesque) and 20 mM Tris-HCl three times, which removed floating particles and contaminants in the particle suspension.

### 4. HS-AFM system

For the HS-AFM measurements, we used a lab-built tip-scan type instrument^33^ with an ultrasmall cantilever (BL-AC10DS-A2, Olympus). The typical resonant frequency and spring constant were 600 kHz and 0.1 N/m in water, respectively. An amorphous carbon tip with a length of ∼500 nm was fabricated on the cantilever bird beak by electron beam deposition. The free oscillation amplitude of the cantilever was ∼5 nm_p-p_ and the setpoint amplitude was approximately 90% of the free amplitude. For the optical image, we first performed the pulling experiments on the particles to search for a microparticle tethered to the substrate via a DNA molecule. Next, the cantilever was maneuvered to approach the surface near the particle. To identify a tethered hairpin construct in the HS-AFM image, the cantilever must be carefully brought close to the particle to avoid attachment of the particle to the cantilever. After the hairpin construct was located in the AFM image, we captured the hairpin dynamics while manipulating the particle with the laser beam controlled by software at the constant velocity of 138 nm/s. After hairpin unfolding was observed, we immediately stopped the laser movement and moved it in the opposite direction with the same velocity.

For kymograph generation, line profiles were manually acquired along the centerline of the DNA frame by frame and were vertically stacked to express the time variation. The apparent size of the globules was expected to correlate with the number of globules. However, the AFM results indicated that the lateral width and the molecular height of the globule varied over time. Therefore, we analyzed the volume of the globule to evaluate its size. For quantitative volume analysis, we first applied a tip deconvolution filter assuming a tip curvature radius of 5 nm^37^. The volume was then calculated by integrating the molecular height in the molecular region with a threshold height of 1 nm from the substrate.

### 5. MD Simulation

We used coarse-grained (CG) models for DNA in the MD simulation because the all-atom (AA) model requires enormous computer resources. For the DNA model, we used 3-Site-Per-Nucleotide (3SPN.2) model^38^, where the individual nucleotides are represented by three CG particles, each representing sugar, phosphate, and nitrogenous base. Although the latest version of the 3SPN model was 3SPN.2C^39^, we used 3SPN.2 because the latter is better tuned to reproduce the physical properties of ssDNA as well as dsDNA^40^. We first built the AA model from the sequence shown in Supplementary Table 2 using X3DNA version 2.4^41^, then converted it to a CG model and performed the simulation using the CafeMol package version 3.2.1 (https://www.cafemol.org)^42^. Langevin dynamics simulations were performed in the NVT ensemble, where the electrolyte temperature was maintained at 298 K using the Nosé–Hoover thermostat and a time step of 0.3 CafeMol time units. The CafeMol time unit was ∼50 fs while effective dynamics were largely accelerated via the use of a low friction coefficient and coarse graining. Each frame was saved at intervals of 300 steps. For the hairpin simulation, the dynamics simulation was performed until unfolding was complete under the condition that the left end of the particle was spatially fixed and the right end of the particle was pulled to the right with a tensile force of 30 pN. Subsequently, the applied force was reduced to 5 pN, and the simulation was performed until refolding was complete.

For generating the pseudo-AFM images, which explicitly consider the existence of the tip, we used a process based on geometric considerations of the mutually excluded volume between the tip and sample^43,44^. For the CG particles of DNA, we used excluded volume diameters of 0.45, 0.64, 0.54, 0.71, 0.49, and 0.64 nm for the phosphate, sugar, adenine base, thymine base, guanine base, and cytosine base, which were taken from the 3SPN.2 model^38^. For the tip, we assumed a sphere with a radius of 20 nm. The topographic images were reconstructed while translating the virtual AFM tip over the surface in XY-Cartesian coordinates at intervals of 0.5 nm/Pixel.

## Acknowledgments

The authors thank Prof. Noriyuki Kodera at Kanazawa University for critical reading and advice on the development of the experimental system. This work was supported by PRESTO, Japan Science and Technology Agency (JST) [JPMJPR20E3 to K.U.]; CREST, JST [JPMJCR13M1 to T.A.]; and KAKENHI, Japan Society for the Promotion of Science [19K15409, 21K04849 (to K.U.), 26119003 and 17H06121 (to T.A.)].

## Author contributions

K.U. and T.A conceived the experimental scheme. K.U. developed the HS-AFM control & analysis software, developed the experimental system, prepared the hairpin concatemer assay, conducted the HS-AFM-OT experiments, and analyzed the data using viscoelastic models and the MD simulation; K.U., S.Y., M.I., S.F., H.W., and T.U. developed the HS-AFM-OT system; K.U. and M.I. designed hairpin synthesis methods; F. N and S.T advised on the simulation; T.A. supervised the whole study. K.U. and T.A. wrote the manuscript.

## References

1. Ashkin, A. & Dziedzic, J. M. Optical trapping and manipulation of viruses and bacteria. Science 235, 1517–1520 (1987).

2. Marago, O. M., Jones, P. H., Gucciardi, P. G., Volpe, G. & Ferrari, A. C. Optical trapping and manipulation of nanostructures. Nat. Nanotechnol. 8, 807–819 (2013).

3. Neuman, K. C. & Nagy, A. Single-molecule force spectroscopy: optical tweezers, magnetic tweezers and atomic force microscopy. Nat. Methods 5, 491–505 (2008).

4. Shrestha, P., Jonchhe, S., Emura, T., Hidaka, K., Endo, M., Sugiyama, H. & Mao, H. B. Confined space facilitates G-quadruplex formation. Nat. Nanotechnol. 12, 582–588 (2017).

5. Shank, E. A., Cecconi, C., Dill, J. W., Marqusee, S. & Bustamante, C. The folding cooperativity of a protein is controlled by its chain topology. Nature 465, 637–640 (2010).

6. Bustamante, C. J., Chemla, Y. R., Liu, S. & Wang, M. D. Optical tweezers in single-molecule biophysics. Nat. Rev. Methods Primers 1, 25 (2021).

7. King, G. A., Gross, P., Bockelmann, U., Modesti, M., Wuite, G. J. L. & Peterman, E. J. G. Revealing the competition between peeled ssDNA, melting bubbles, and S-DNA during DNA overstretching using fluorescence microscopy. Proc. Natl. Acad. Sci. USA 110, 3859–3864 (2013).

8. Zhang, X. H., Qu, Y. Y., Chen, H., Rouzina, I., Zhang, S. L., Doyle, P. S. & Yan, J. Interconversion between Three Overstretched DNA Structures. J. Am. Chem. Soc. 136, 16073–16080 (2014).

9. Neupane, K., Foster, D. A. N., Dee, D. R., Yu, H., Wang, F. & Woodside, M. T. Direct observation of transition paths during the folding of proteins and nucleic acids. Science 352, 239–242 (2016).

10. Cheng, W., Arunajadai, S. G., Moffitt, J. R., Tinoco, I. & Bustamante, C. Single-Base Pair Unwinding and Asynchronous RNA Release by the Hepatitis C Virus NS3 Helicase. Science 333, 1746–1749 (2011).

11. Huguet, J. M., Bizarro, C. V., Forns, N., Smith, S. B., Bustamante, C. & Ritort, F. Single-molecule derivation of salt dependent base-pair free energies in DNA. Proc. Natl. Acad. Sci. USA 107, 15431–15436 (2010).

12. Woodside, M. T., Behnke-Parks, W. M., Larizadeh, K., Travers, K., Herschlag, D. & Block, S. M. Nanomechanical measurements of the sequence-dependent folding landscapes of single nucleic acid hairpins. Proc. Natl. Acad. Sci. USA 103, 6190–6195 (2006).

13. Fazal, F. M., Meng, C. A., Murakami, K., Kornberg, R. D. & Block, S. M. Real-time observation of the initiation of RNA polymerase II transcription. Nature 525, 274–277 (2015).

14. Colom, A., Derivery, E., Soleimanpour, S., Tomba, C., Dal Molin, M., Sakai, N., Gonzalez-Gaitan, M., Matile, S. & Roux, A. A fluorescent membrane tension probe. Nat. Chem. 10, 1118–1125 (2018).

15. Bornschlögl, T., Romero, S., Vestergaard, C. L., Joanny, J. F., Nhieu, G. T. V. & Bassereau, P. Filopodial retraction force is generated by cortical actin dynamics and controlled by reversible tethering at the tip. Proc. Natl. Acad. Sci. USA 110, 18928–18933 (2013).

16. Leijnse, N., Oddershede, L. B. & Bendix, P. M. Helical buckling of actin inside filopodia generates traction. Proc. Natl. Acad. Sci. USA 112, 136–141 (2015).

17. Heller, I., Sitters, G., Broekmans, O. D., Farge, G., Menges, C., Wende, W., Hell, S. W., Peterman, E. J. G. & Wuite, G. J. L. STED nanoscopy combined with optical tweezers reveals protein dynamics on densely covered DNA. Nat. Methods 10, 910–916 (2013).

18. Diekmann, R., Wolfson, D. L., Spahn, C., Heilemann, M., Schuttpelz, M. & Huser, T. Nanoscopy of bacterial cells immobilized by holographic optical tweezers. Nat. Commun. 7, 13711 (2016).

19. Kodera, N., Yamamoto, D., Ishikawa, R. & Ando, T. Video imaging of walking myosin V by high-speed atomic force microscopy. Nature 468, 72 (2010).

20. Shibata, M., Yamashita, H., Uchihashi, T., Kandori, H. & Ando, T. High-speed atomic force microscopy shows dynamic molecular processes in photoactivated bacteriorhodopsin. Nat. Nanotechnol. 5, 208–212 (2010).

21. Ando, T., Uchihashi, T. & Scheuring, S. Filming Biomolecular Processes by High-Speed Atomic Force Microscopy. Chem. Rev. 114, 3120–3188 (2014).

22. Ando, T. High-speed atomic force microscopy. Curr. Opin. Chem. Biol. 51, 105–112 (2019).

23. Kodera, N., Noshiro, D., Dora, S. K., Mori, T., Habchi, J., Blocquel, D., Gruet, A., Dosnon, M., Salladini, E., Bignon, C., Fujioka, Y., Oda, T., Noda, N. N., Sato, M., Lotti, M., Mizuguchi, M., Longhi, S. & Ando, T. Structural and dynamics analysis of intrinsically disordered proteins by high-speed atomic force microscopy. Nat. Nanotechnol. 16, 181–189 (2021).

24. Heath, G. R. & Scheuring, S. Advances in high-speed atomic force microscopy (HS-AFM) reveal dynamics of transmembrane channels and transporters. Curr. Opin. Struct. Biol. 57, 93–102 (2019).

25. Wickham, S. F. J., Endo, M., Katsuda, Y., Hidaka, K., Bath, J., Sugiyama, H. & Turberfield, A. J. Direct observation of stepwise movement of a synthetic molecular transporter. Nat. Nanotechnol. 6, 166–169 (2011).

26. Nievergelt, A. P., Banterle, N., Andany, S. H., Gonczy, P. & Fantner, G. E. High-speed photothermal off-resonance atomic force microscopy reveals assembly routes of centriolar scaffold protein SAS-6. Nat. Nanotechnol. 13, 696–701 (2018).

27. Umeda, K., McArthur, S. J. & Kodera, N. Spatiotemporal resolution in high-speed atomic force microscopy for studying biological macromolecules in action. Microscopy 72, 151–161 (2023).

28. Umeda, K., Okamoto, C., Shimizu, M., Watanabe, S., Ando, T. & Kodera, N. Architecture of zero-latency ultrafast amplitude detector for high-speed atomic force microscopy. Appl. Phys. Lett. 119, 181602 (2021).

29. Lin, Y. C., Guo, Y. R., Miyagi, A., Levring, J., MacKinnon, R. & Scheuring, S. Force-induced conformational changes in PIEZO1. Nature 573, 230–234 (2019).

30. Yu, J., Zhang, Z. F., Cao, K. & Huang, X. T. Visualization of alkali-denatured supercoiled plasmid DNA by atomic force microscopy. Biochem. Biophys. Res. Commun. 374, 415–418 (2008).

31. Maaloum, M., Beker, A. F. & Muller, P. Secondary structure of double-stranded DNA under stretching: Elucidation of the stretched form. Phys. Rev. E 83, 031903 (2011).

32. Martonfalvi, Z. & Kellermayer, M. Individual Globular Domains and Domain Unfolding Visualized in Overstretched Titin Molecules with Atomic Force Microscopy. PLoS ONE 9, e85847 (2014).

33. Fukuda, S., Uchihashi, T., Iino, R., Okazaki, Y., Yoshida, M., Igarashi, K. & Ando, T. High-speed atomic force microscope combined with single-molecule fluorescence microscope. Rev. Sci. Instrum. 84, 073706 (2013).

34. Bikard, D., Loot, C., Baharoglu, Z. & Mazel, D. Folded DNA in Action: Hairpin Formation and Biological Functions in Prokaryotes. Microbiol. Mol. Biol. Rev. 74, 570–588 (2010).

35. Neupane, K., Ritchie, D. B., Yu, H., Foster, D. A. N., Wang, F. & Woodside, M. T. Transition Path Times for Nucleic Acid Folding Determined from Energy-Landscape Analysis of Single-Molecule Trajectories. Phys. Rev. Lett. 109, 068102 (2012).

36. Hoffer, N. Q., Neupane, K., Pyo, A. G. T. & Woodside, M. T. Measuring the average shape of transition paths during the folding of a single biological molecule. Proc. Natl. Acad. Sci. USA 116, 8125–8130 (2019).

37. Villarrubia, J. S. Algorithms for scanned probe microscope image simulation, surface reconstruction, and tip estimation. J. Res. Natl. Inst. Stand. Technol. 102, 425–454 (1997).

38. Hinckley, D. M., Freeman, G. S., Whitmer, J. K. & de Pablo, J. J. An experimentally-informed coarse-grained 3-site-per-nucleotide model of DNA: Structure, thermodynamics, and dynamics of hybridization. J. Chem. Phys. 139, 144903 (2013).

39. Freeman, G. S., Hinckley, D. M., Lequieu, J. P., Whitmer, J. K. & de Pablo, J. J. Coarse-grained modeling of DNA curvature. J. Chem. Phys. 141, 165103 (2014).

40. Chakraborty, D., Horio, N. & Thirumalai, D. Sequence-Dependent Three Interaction Site Model for Single- and Double-Stranded DNA. J. Chem. Theory Comput. 14, 3763–3779 (2018).

41. Lu, X. J. & Olson, W. K. 3DNA: a versatile, integrated software system for the analysis, rebuilding and visualization of three-dimensional nucleic-acid structures. Nat. Protoc. 3, 1213–1227 (2008).

42. Kenzaki, H., Koga, N., Hori, N., Kanada, R., Li, W. F., Okazaki, K., Yao, X. Q. & Takada, S. CafeMol: A Coarse-Grained Biomolecular Simulator for Simulating Proteins at Work. J. Chem. Theory Comput. 7, 1979–1989 (2011).

43. Markiewicz, P. & Goh, M. C. Simulation of atomic force microscope tip–sample/sample–tip reconstruction. J. Vac. Sci. Technol. B 13, 1115–1118 (1995).

44. Ando, T., Kodera, N., Uchihashi, T., Miyagi, A., Nakakita, R., Yamashita, H. & Matada, K. High-speed Atomic Force Microscopy for Capturing Dynamic Behavior of Protein Molecules at Work. e-J. Surf. Sci. Nanotechnol. 3, 384–392 (2005).

